# An optimized messenger RNA vaccine candidate protects non-human primates from Zika virus infection

**DOI:** 10.1101/2022.10.11.511814

**Authors:** Brooke Bollman, Naveen Nunna, Kapil Bahl, Chiaowen Joyce Hsiao, Hamilton Bennett, Scott Butler, Bryant Foreman, Katherine E. Burgomaster, Maya Aleshnick, Wing-Pui Kong, Brian E. Fisher, Tracy J. Ruckwardt, Kaitlyn M. Morabito, Barney S. Graham, Kimberly A. Dowd, Theodore C. Pierson, Andrea Carfi

**Author notes:** **Corresponding author:** Brooke Bollman, PhD, Moderna, Inc., 200 Technology Square, Cambridge, MA, 02139 USA.

## Abstract

Zika virus (ZIKV), an arbovirus transmitted by mosquitoes, was identified as a cause of congenital disease during a major outbreak in the Americas in 2016. Vaccine design strategies relied on limited available isolate sequence information due to the rapid response necessary. The first-generation ZIKV mRNA vaccine, mRNA-1325, was initially generated and, as additional strain sequences became available, a second mRNA vaccine, mRNA-1893, was developed. Herein, we compared the immune responses following mRNA-1325 and mRNA-1893 vaccination and reported that mRNA-1893 generated comparable neutralizing antibody titers to mRNA-1325 at 1/20^th^ of the dose and provided complete protection from ZIKV challenge in non-human primates. In depth characterization of these vaccines indicated that the observed immunologic differences could be attributed to a single amino acid residue difference that compromised mRNA-1325 virus-like particle formation.

## Introduction

Zika virus (ZIKV), an arbovirus transmitted by mosquitoes, is a member of the *Flaviviridae* family, which includes yellow fever, dengue, and West Nile viruses^1^. ZIKV was first identified in 1947, and, until 2007, caused sporadic infections across Africa. In 2007, a large-scale epidemic occurred on Yap Island, followed by a larger outbreak in French Polynesia in 2013^2^. Complications associated with ZIKV infection during pregnancy, including microcephaly and other congenital abnormalities in the developing fetus and newborn^3^, were reported during the 2015 outbreak of ZIKV in the Western Hemisphere and led the World Health Organization to declare ZIKV a Public Health Emergency of International Concern in 2016^4^. In addition to maternal–fetal transmission^5–7^, sexual transmission of ZIKV has been reported^8^. Based upon prior genetic and phylogenetic analyses, 2 ZIKV lineages have been identified and classified as African and Asian, with the latter traced to recent outbreaks^9–11^.

ZIKV contains an 11-kilobase, non-segmented, positive-sense, single-stranded RNA genome. The genome contains 5′ and 3′ untranslated regions flanking a single open reading frame (ORF) that encodes a single polyprotein that is cleaved into 3 structural proteins (capsid [C], premembrane/membrane [prM], and envelope [E]) and 7 non-structural proteins (NS1, NS2A, NS2B, NS3, NS4A, NS4B, and NS5)^12^. Within the secretory pathway, prM is cleaved by the host furin protease during maturation to produce M and pr. When the virion exits the cell and enters the neutral pH of the extracellular environment, pr is released from the virion^13,14^. E is the major antigenic determinant of the virus and mediates receptor binding and virus–cell membrane fusion during entry^15,16^. Since E is the primary target for neutralizing antibodies (nAbs), it is the antigen targeted by most vaccines^17^. Expression of prME in mammalian cells produces non-infectious virus-like particles (VLPs) that share antigenic features with infectious virions^18,19^; VLPs retain key properties of viruses, including fusogenic activity^20^ and the induction of a nAb response.

As the ZIKV epidemic required a rapid public health response, vaccine design strategies relied on the limited isolate sequence information available at the time. The public bank of full-length human ZIKV sequences contained 8 sequences in 2014; this increased to 30 by March 2016^21^. The first-generation mRNA vaccine, mRNA-1325 (Moderna, Inc., Cambridge, MA), encodes the prME ORF from a Micronesia 2007 ZIKV isolate^22^, with an upstream immunoglobulin E (IgE) leader sequence to drive secretion. As additional sequences became available, a second mRNA vaccine was generated, mRNA-1893, which encodes the contemporary RIO-U1 Zika prME ORF. RIO-U1 was isolated from the urine of a pregnant woman with symptoms of an acute phase ZIKV infection^21,23^. mRNA-1893 encodes a Japanese encephalitis virus (JEV)–derived leader sequence, which has been reported to improve host signalase cleavage^18,24^. Aside from the different leader sequence, mRNA-1325 and mRNA-1893 have 5 amino acid residue differences in the prME ORF (2 in the pr region; 3 in the E gene).

Here, we compared mRNA-1325 and mRNA-1893 and showed that mRNA-1893 elicited similar nAb levels at lower doses compared with mRNA-1325 and provided complete protection from ZIKV challenge in non-human primates (NHPs). Through further characterization of mRNA-1893 and mRNA-1325, we identified a single amino acid residue difference that compromises mRNA-1325 VLP formation and may underlie the immunologic differences observed between the vaccines.

## Results

### mRNA-1893 generates VLPs in vitro

mRNA-1893 and mRNA-1325 were formulated in ionizable lipid based LNPs and tested for intracellular expression levels of prME and differences in VLP production following in vitro transfection. The antibody ZV-116 was used to detect E protein in cell lysates and VLPs purified from supernatant^25^. E protein was detected in cell lysates from mRNA-1325 and mRNA-1893, and in mRNA-1893 supernatant (**Figure 1a**). Bands corresponding to the approximate molecular weight of uncleaved prM were detected for mRNA-1325 and mRNA-1893 but were more prominent for mRNA-1893 (**Figure 1b**). In addition, prM was not detected in supernatant, but a band of the approximate size of cleaved M protein was present in mRNA-1893 supernatant, suggesting the production of mature VLPs. An additional band above 50 kD was present in all cell lysates, including the control, reflecting a non-specific band.

**Figure 1.**
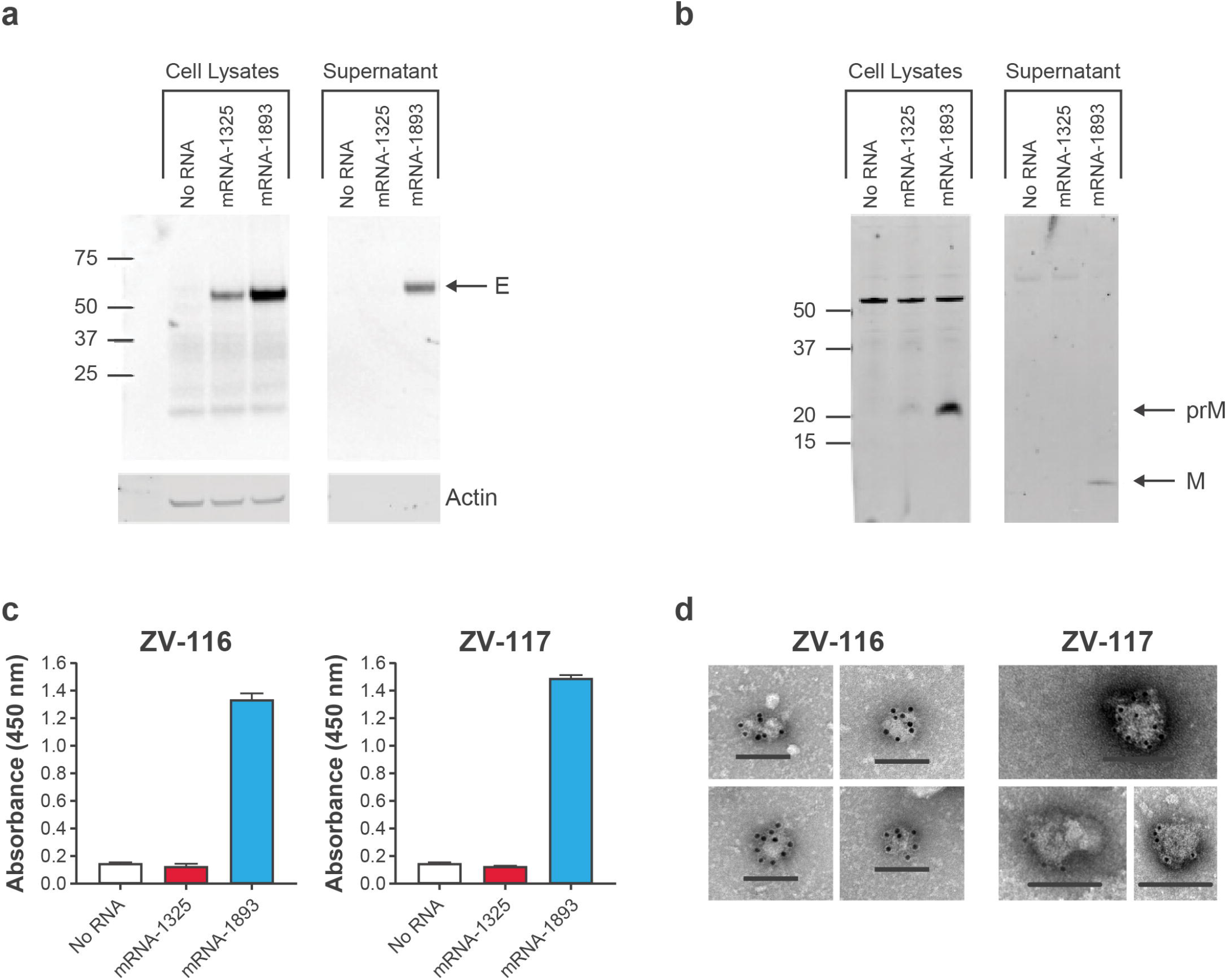
VLP generation from mRNA-1893 in vitro. (**a**) Cell lysates and VLP pellets were blotted for E protein using ZV-116 antibody. Actin was used as a loading control for cell lysate samples. (**b**) Cell lysates and VLP pellets were blotted for prM and M proteins using an anti-Zika prM polyclonal antibody. (**c**) VLP secretion was measured by sandwich ELISA. Data are representative of a single experiment, with error bars indicating the SD of technical repeats on the same plate (ie, triplicate wells). (**d**) For negative staining and immunogold labeling, ZV-116 at 30,000X magnification (left) or ZV-117 at 50,000X magnification (right) was used, followed by the use of a gold-labeled anti-human secondary antibody. Scale bar indicates 100 nm. E, envelope; ELISA, enzyme-linked immunosorbent assay; M, membrane; prM, premembrane/membrane; SD, standard deviation; VLP, virus-like particle.

To differentiate between monomeric E protein and assembled VLPs, a sandwich enzyme-linked immunosorbent assay (ELISA) was performed using the anti-E antibodies ZV-116 or ZV-117 (for capture) and ZV-67 (for detection). Both ZV-116 and ZV-67 recognize a linear epitope in the Domain III lateral ridge^26^ and recognize free E monomers or VLP-associated E protein. ZV-117 binds across an E protein dimer and preferentially recognizes E protein incorporated into VLPs, although it can also weakly bind to E protein monomer^27^. Supernatants collected 48 hours after transfection were added to plates coated with either ZV-116 or ZV-117 (**Figure 1c**). E protein was detected in supernatant following mRNA-1893 transfection using ZV-116 or ZV-117. However, levels of supernatant E protein following mRNA-1325 transfection were comparable to the untransfected controls with both antibodies, suggesting that mRNA-1325 does not generate secreted VLPs in vitro.

Transmission electron microscopy and negative staining, used to characterize VLP composition generated after mRNA-1893 transfection, revealed that VLPs had a roughly spherical structure, with an average diameter of 50 to 100 nm. Particles were imaged by immunogold labeling using ZV-116 and ZV-117 antibodies. VLPs were reactive with both antibodies (**Figure 1d**), consistent with the ELISA results. Collectively, the data demonstrate that mRNA-1893 transfection in vitro results in the formation and secretion of VLPs.

### Intracellular E protein generated by mRNA-1325 has a short half-life and is rapidly degraded by the proteasome

In vitro experiments were conducted to examine the apparent absence of VLPs or E protein dimers after mRNA-1325 transfection despite detectable E protein expression, albeit at lower levels than those seen for mRNA-1893 (**Figure 1a**). As rapid protein degradation could contribute to the lack of VLP generation, protein half-life was examined. Intracellular E protein was detected by ZV-116 (**Figure 2a**) and a pan-flavivirus antibody 4G2 (**Figure 2b**)^16,28^. While mRNA-1893 protein had a half-life of ≥ 6 hours, mRNA-1325 had a relatively short half-life of <2 hours (**Figure 2a**). In addition, no E protein was detected at any time after transfection for mRNA-1325 using 4G2, suggesting that protein produced by mRNA-1325 could be misfolded (**Figure 2b**). To further elucidate the pathway involved in the accelerated E protein degradation, 293T cells transfected with mRNA-1325 were treated with cycloheximide (CHX) plus chloroquine or CHX plus MG132 to inhibit lysosomal proteolysis or ubiquitin-mediated proteasomal degradation pathways, respectively. Chloroquine had no effect on E protein half-life (**Figure 2c**), whereas MG132 extended E protein half-life ≥ 8 hours after transfection (**Figure 2d**). These results suggest that mRNA-1325–expressed E protein is at least partially misfolded and undergoes rapid proteasomal degradation, thus leading to loss of VLP production.

**Figure 2.**
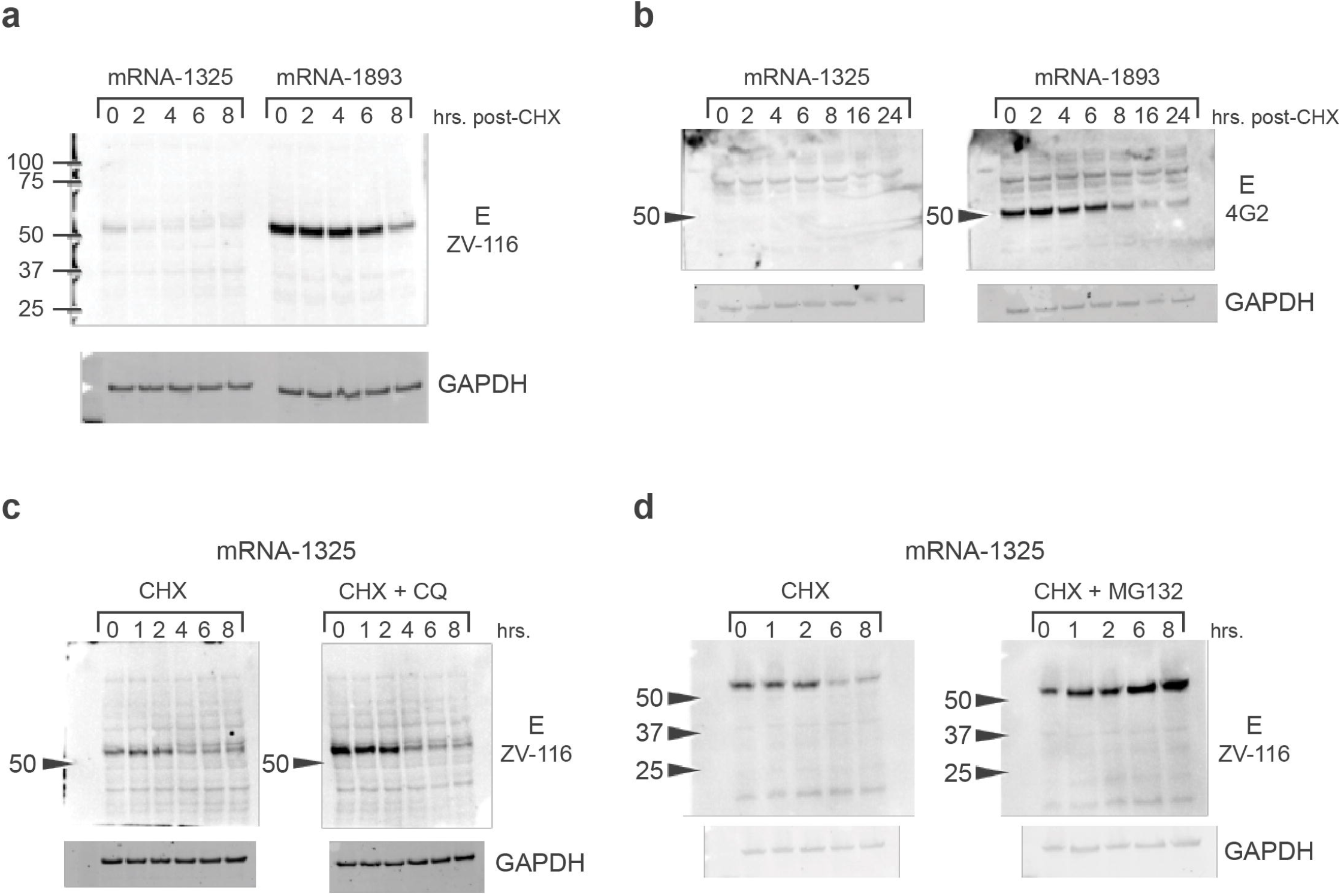
Time course of intracellular E protein generated by mRNA-1325. (**a**) E protein expression was assessed in cell lysates from mRNA-transfected cells, following the addition of CHX and using antibody ZV-116. (**b**) E protein levels were also assayed, following the addition of CHX and using the anti-flavivirus fusion loop antibody 4G2, under non-reducing conditions. (**c**) 293T cells were transfected with mRNA-1325 and were treated with CHX or CHX + CQ, harvested at the indicated time points following treatment, and blotted for E using ZV-116. (**d**) 293T cells were transfected with mRNA-1325 and were treated with CHX or CHX + MG132 to block proteasomal degradation. CHX, cycloheximide; CQ, chloroquine; E, envelope.

### Alanine to threonine substitution at residue 40 of the E protein in mRNA-1325 rescues VLP production in vitro and enhances nAb titers in mice

mRNA-1325 and mRNA-1893 differ in signal peptides (IgE and JEV, respectively) and in 5 amino acid residues (valine, serine, alanine, valine, and threonine in mRNA-1325 substituted for alanine, asparagine, threonine, methionine, and methionine in mRNA-1893, respectively) in the prME ORF (**Figure 3a**). To investigate whether these differences contributed to the VLP production defect of mRNA-1325, a panel of mRNA-1325 sequence variants was generated.

**Figure 3.**
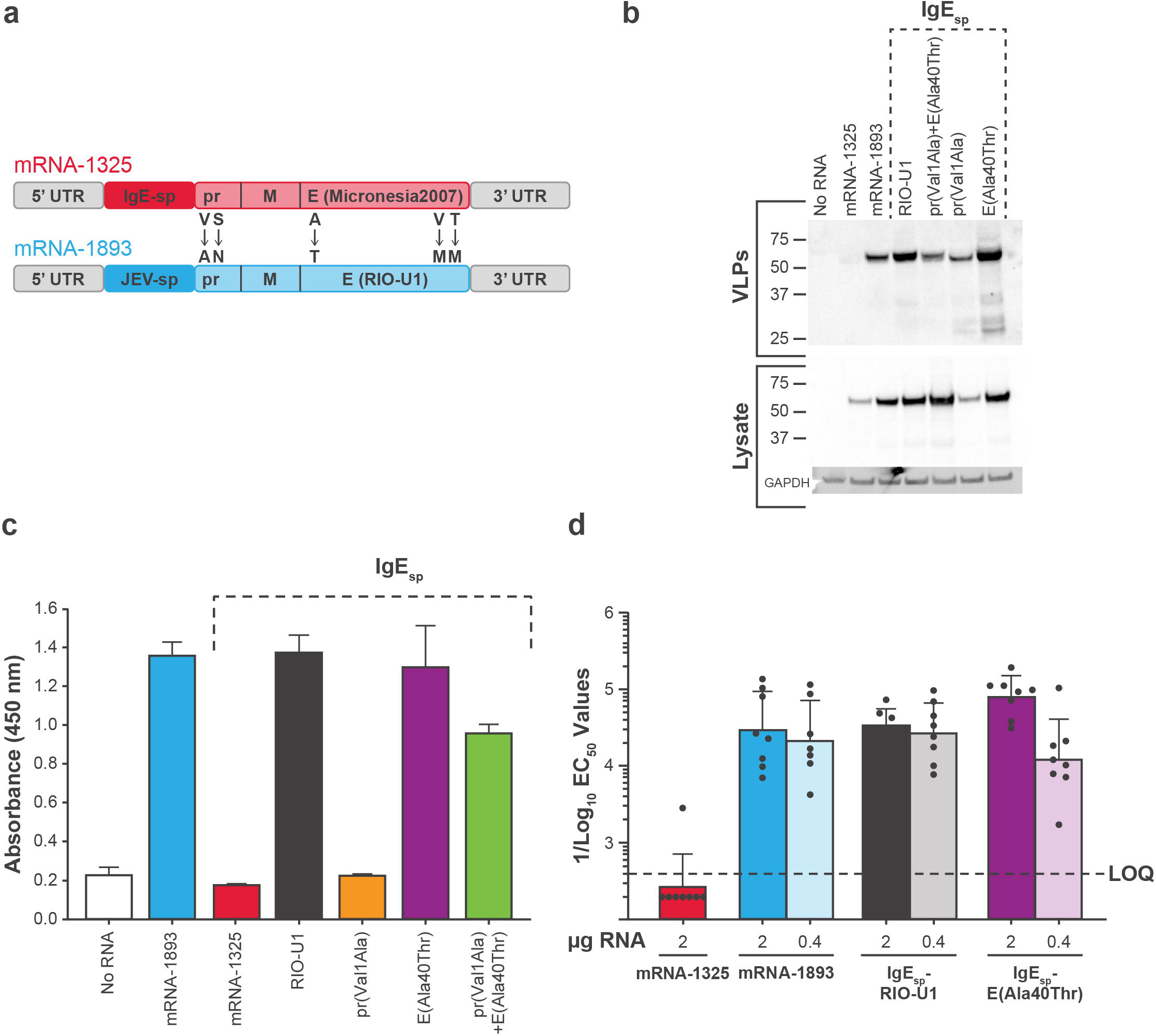
VLP formation for an alanine to threonine substitution in the mRNA-1325 E ORF. (**a**) mRNA-1325 and mRNA-1893 differ in signal sequences and 5 amino acid residues within the prME ORF. (**b**) Cell lysates and VLP pellets were blotted for E protein using ZV-116 antibody. GAPDH was used as a loading control. (**c**) VLP secretion from transfected cells was measured in a sandwich ELISA with ZV-117 antibody, followed by ZV-67 antibody, and an anti-mouse HRP-conjugated secondary antibody. (**d**) Colored bars represent GMT for the group, and black circles represent individual animals (n = 8 per group). Error bars indicate SD from the GMT. Dashed line indicates assay LOQ. E, envelope; EC_50_, half-maximal effective concentration; ELISA, enzyme-linked immunosorbent assay; GMT, geometric mean titer; HRP, horseradish peroxidase; IgE, immunoglobulin E; JEV, Japanese encephalitis virus; LOQ, limit of quantitation; ORF, open reading frame; prME, premembrane/membrane and envelope; RVP, reporter virus particle; SD, standard deviation; UTR, untranslated region; VLP, virus-like particle; ZIKV, Zika virus.

Due to their proximity to the amino terminus and potential to affect VLP production, the residues pr1 and E40 were targeted to generate the following variants: IgE_sp_-pr(Val1Ala); IgE_sp_-E(Ala40Thr); and IgE_sp_-pr(Val1Ala) + E(Ala40Thr). To exclude the possibility of the leader sequence affecting VLP production, IgE_sp_-RIO-U1 was generated, which contains the entire prME ORF from mRNA-1893 but retains the IgE_sp_ from mRNA-1325.

mRNA-1325, mRNA-1893, and the variants were transiently transfected into 293T cells, and lysates and supernatant were assayed for E protein expression and VLP production. All variants rescued VLP production to levels detectable by Western blot (**Figure 3b**). A sandwich ELISA using ZV-117 and ZV-67 antibodies demonstrated that mRNA-1893, IgE_sp_-RIO-U1, and IgE_sp_-E(Ala40Thr) produced similar levels of secreted VLPs (**Figure 3c**).

To determine whether the rescue of VLP production in vitro was correlated with an increase in immunogenicity in vivo, C57BL/6 mice were immunized intramuscularly at days 1 and 29 with 2 μg or 0.4 μg LNPs containing mRNA-1325, mRNA-1893, or variants. Seven of 8 mice immunized with mRNA-1325 2 μg had nAb titers below the assay detection limit. Conversely, animals immunized with mRNA-1893 2 μg or 0.4 μg or sequence variants IgE_sp_-RIO-U1 or IgE_sp_-E(Ala40Thr) had half-maximal effective concentration (EC_50_) values of >1/10^3^, with most animals having titers between 1/10^4^ and 1/10^5^ (**Figure 3d**). These data indicated that the VLP production defects and lower immunogenicity observed with mRNA-1325 are not due to the IgE_sp_ leader sequence, but instead can be mapped to the presence of alanine at E40 instead of threonine.

### mRNA-1893 shows superior immunogenicity and efficacy compared with mRNA-1325 in rhesus macaques

To evaluate the immunogenicity and efficacy of mRNA-1325 and mRNA-1893 in rhesus macaques, both mRNA constructs were formulated in ionizable lipid based LNPs and administered intramuscularly using a 2-dose regimen (days 1, 29). mRNA-1325 was tested at 3 concentrations (10, 50, and 200 μg) and mRNA-1893 was tested at the 10-μg concentration. A non-coding mRNA control (200 μg) and mRNA-1325 delivered as single 200-μg dose (day 1) were also included. nAbs were measured using the ZIKV RVP neutralization assay^29^ at week 4 before boost and immediately before challenge at week 8.

Among mRNA-1325 regimens at week 4, a significant dose response was observed for the 2-dose, 10-μg compared with the 1-dose, 200-μg regimen, but not with the 2-dose, 200-μg regimen (*P*<0.05 and *P*>0.05, respectively; **Figure 4a; Supplementary Table 1**). At week 8, a dose response was observed for the low-vs high-dose 2-dose regimens of mRNA-1325 (10 μg vs 200 μg; *P*<0.001); mean nAb titers for the 10-, 50-, and 200-μg doses were 1697, 4347, and 7659, respectively. When comparing the 1-vs 2-dose regimens at week 8, mRNA-1325 200 μg generated significantly higher nAb titers when administered as a 2-dose compared with a 1-dose regimen (*P*<0.05). Comparisons between constructs at the same dose at week 8 showed a significantly lower nAb response for mRNA-1325 compared with mRNA-1893 when the vaccines were administered as a 10-μg, 2-dose regimens (*P*<0.05). Surprisingly, nAb titers in mRNA-1893 10-μg, 2-dose regimen recipients were comparable to titers in mRNA-1325 200-μg, 2-dose regimen recipients (6479 vs 7659, respectively; *P*≈0.97), suggesting that mRNA-1893 generated similar levels of nAbs at 1/20^th^ of the dose and was consistent with the increase in VLP formation observed in the in vitro experiments.

**Figure 4.**
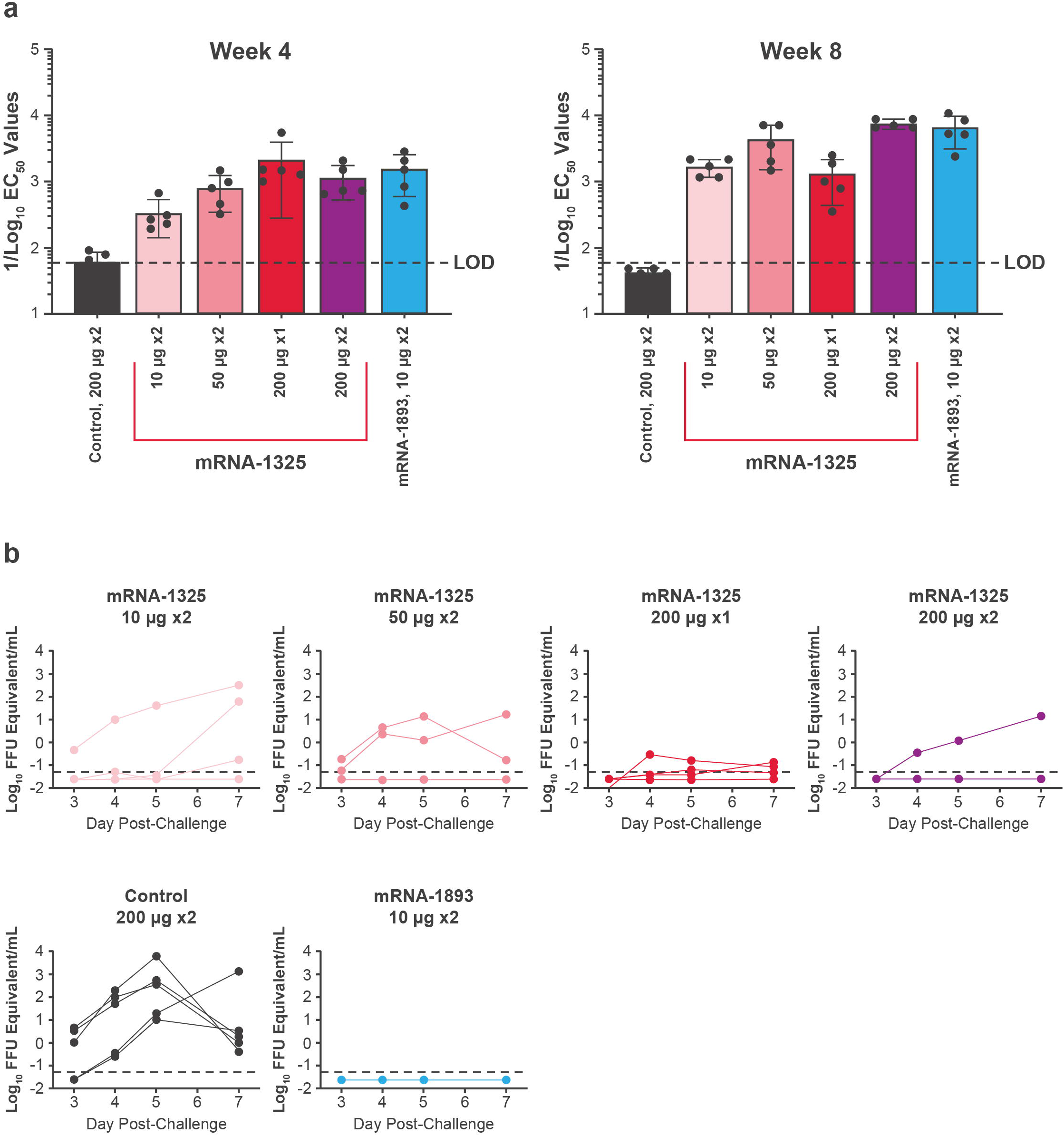
nAb titers and protection of mRNA-1325 and mRNA-1893 against challenge in NHPs. (**a**) nAb titers were measured using the ZIKV RVP assay on week 4 (day 28) and week 8 (day 56). Colored bars represent GMTs for each group (mean ± SD) and black circles represent individual animals (n = 5 per group). Dashed line indicates assay LOD. (**b**) Viremia was detected via qPCR on days 3, 4, 5, and 7 following challenge. Dashed line indicates assay LOD. EC_50_ half-maximal effective concentration; FFU, focus forming units; GMT, geometric mean titer; LOD, limit of detection; nAb, neutralizing antibody; NHP, non-human primate; qPCR, quantitative polymerase chain reaction; RVP, reporter virus particle; VLP, virus-like particle; ZIKV, Zika virus.

All groups were challenged with 1 × 10^3^ plaque-forming units ZIKV (strain PRVABC59) delivered subcutaneously on day 56 and assayed for viremia by quantitative reverse transcription polymerase chain reaction on days 3, 4, 5, and 7 thereafter. All animals in the non-coding mRNA control group and ≥ 1 animal from each of the mRNA-1325 groups had detectable viremia (**Figure 4b**). No viremia was detected in animals in the mRNA-1893 group following challenge.

To evaluate whether mRNA-1893 vaccination generated sterilizing immunity, nAb titers were evaluated 14 and 28 days following challenge for individual animals in each dose group (**Figure 5**). EC_50_ values for ≥ 1 animal in each of the mRNA-1325 groups had a ≥ 4-fold anamnestic response. All animals that had a ≥4-fold increase also had detectable viremia following challenge (**Figure 5**). No animals vaccinated with mRNA-1893 had a 4-fold increase in neutralization activity samples analyzed post-challenge. Thus, immune responses elicited by mRNA-1893 rapidly control infection, thereby preventing an anamnestic response. Collectively, these data indicate that mRNA-1893 generates more robust protection at doses substantially lower than mRNA-1325 in NHPs.

**Figure 5.**
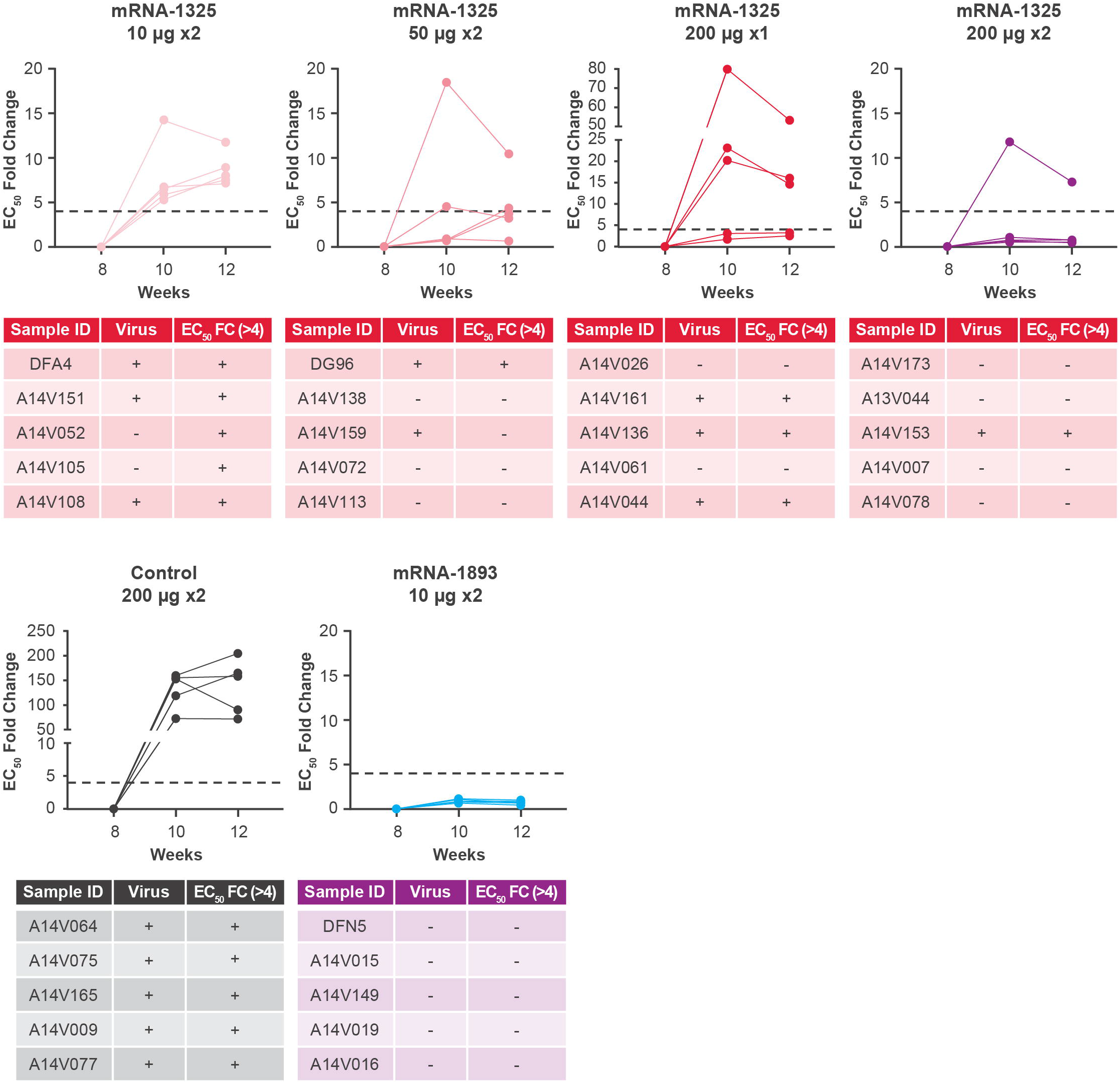
nAb titers in NHPs immunized with mRNA-1893. Data is plotted as the EC_50_ fold change relative to time of challenge (week 8), which was set to 1. Tables below each graph indicate whether the virus was detected on any day following challenge and if the EC_50_ value had a 4-fold or greater increase following challenge. n = 5 per group. EC_50_ half-maximal effective concentration; FC, fold change; nAb, neutralizing antibody; NHP, non-human primate; RVP, reporter virus particle; ZIKV, Zika virus.

### ZIKV CprME–based mRNA constructs produce C-containing VLPs in vitro and are immunogenic in mice

The ZIKV ORF encodes the structural proteins C, prM, and E^30–32^. Expression of prME is sufficient for VLP formation, but it is possible that C-containing VLPs (CprME) are more immunogenic than prME-only VLPs^32–34^. Two CprME-encoding constructs were generated, including 1 encoding wild-type RIO-U1 Zika CprME ORF, and compared with mRNA-1893, both for VLP production in vitro and nAb responses in mice. As production of CprME VLPs requires C-prM cleavage by NS2B3 protease^30–32^, we generated a second mRNA encoding RIO-U1 NS2B3. As an alternative method of generating cleaved C, a C-2A-prME construct was designed with a ribosome 2A stop-go sequence between C and prM. Insertion of this sequence should result in release of C from the nascent polyprotein during translation, eliminating the need for an NS2B3-mediated cleavage step^35^.

The mRNA constructs encoding wild-type CprME, either alone or with NS2B3 mRNA, and C-2A-prME (alone) were transfected into 293T cells side-by-side with mRNA-1893 and variants IgE_sp_-RIO-U1 and IgE_sp_-E(Ala40Thr). VLP production was compared across constructs by purifying VLPs from supernatant and Western blotting for E and prM proteins (**Figure 6a, 6b**). Both C-containing constructs produced VLPs, though as expected, the CprME construct required the co-transfection of NS2B3. The CprME/NS2B3 combination resulted in a higher amount of detectable E protein than any of the prME-based constructs or C-2A-prME. In addition, the presence of the low molecular weight band representing cleaved M protein indicates that the VLPs produced by these C constructs are mature (**Figure 6a**). To investigate whether the VLPs did in fact contain C, cell lysates and purified VLPs from transfected cells were blotted for the C protein (**Figure 6b**). The C protein was detected in the lysates of both constructs, with C-2A-prME producing a slightly higher molecular weight protein due to the 19-amino acid residue tail generated by the 2A sequence. The C protein was also detected in the purified VLPs from CprME/NS2B3 co-transfection, but not transfection of C-2A-prME. As the E protein Western blot on purified VLPs indicates that the C-2A-prME construct produces VLPs, the lack of signal could be due to an overall lower level of expression and VLP production of C-2A-prME compared with CprME/NS2B3, or to the lack of packaging of C protein, resulting in prME-only VLPs (**Figure 6a**).

**Figure 6.**
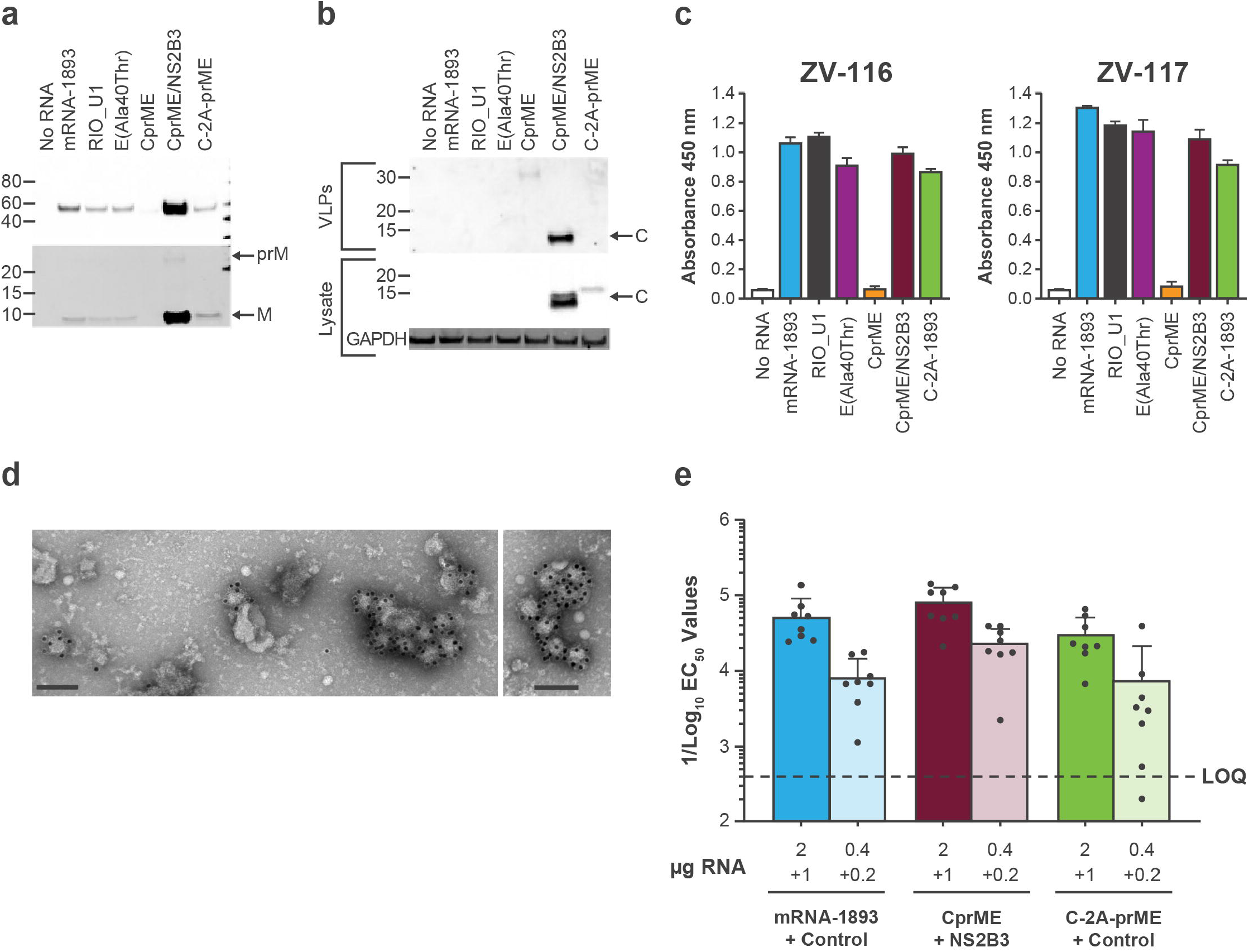
ZIKV CprME mRNA VLP generation in vitro and immunogenicity in mice. (**a**) The VLP pellet was blotted for E protein using ZV-116 (top) and prM and M proteins were blotted using an anti-Zika prM polyclonal antibody (bottom). Arrows indicate prM and M. (**b**) Purified VLPs (top) and lysates from transfected cells (bottom) were blotted for C protein using an anti-Zika capsid polyclonal antibody. (**c**) VLP secretion was measured by sandwich ELISA, with either ZV-116 (left) or ZV-117 (right), followed by the addition of the anti-E antibody ZV-67 and an anti-mouse HRP-conjugated secondary antibody. Data are representative of a single experiment, with error bars indicating the SD of technical repeats on the same plate (ie, triplicate wells). (**d**) Cells were transfected with CprME + NS2B3 mRNAs. Concentrated VLPs were stained with uranyl acetate and labeled with ZV-116 at 30,000X magnification, followed by a gold-labeled anti-human secondary antibody. Scale bar indicates 100 nm. (**e**) nAb titers were measured using the ZIKV RVP neutralization assay on day 56 following 2 doses of mRNA administered on day 1 and day 29. Colored bars represent GMT for each group (mean ± SD), and black circles represent individual animals (n = 8 per group). Dashed line indicates assay LOQ. C, capsid; CprME, capsid, premembrane/membrane, and envelope; E, envelope; EC_50_ half-maximal effective concentration; ELISA, enzyme-linked immunosorbent assay; GMT, geometric mean titer; HRP, horseradish peroxidase; LOQ, limit of quantitation; M, membrane; nAb, neutralizing antibody; prM, premembrane/membrane; prME, premembrane/membrane and envelope; RVP, reporter virus particle; SD, standard deviation; VLP, virus-like particle; ZIKV, Zika virus.

As C protein stability is important for the assembly of infectious flavivirus VLPs^36^, the inclusion of C may allow CprME VLPs to better withstand the purification process than prME-only particles, and therefore, give the appearance of increased VLP production. For more accurate comparisons of VLP production levels across constructs, supernatant from transfected cells was used directly in the VLP sandwich ELISAs (**Figure 6c**). Both ZV-116 and ZV-117 ELISAs detected supernatant E protein from transfection of CprME/NS2B3 and C-2A-prME, but not CprME alone. VLPs were detected for both C-containing designs and prME-only constructs. VLP production was further confirmed by the Western blot of purified VLPs (**Figure 6a**).

Electron microscopy confirmed the presence of VLPs (**Figure 6d**). The quantities of CprME/NS2B3 and prME VLPs varied between the ELISA and Western blot results; while this may indicate differential stability between the VLPs, we do not exclude the possibility of this variation arising from differing purification conditions.

To determine whether the CprME VLPs were better immunogens than prME-only VLPs, C57BL/6 mice were immunized intramuscularly with LNPs containing mRNA-1893/non-coding control, CprME/NS2B3, or C-2A-prME/non-coding control. Each combination contained a mass ratio of 2 parts structural protein mRNA to 1-part non-coding or NS2B3 mRNA; 3 or 0.6 μg of total mRNA were administered. Animals received 2 doses 28 days apart and were phlebotomized at day 56 for serum nAb analysis using ZIKV RVPs. All constructs at the high dose (3 μg) generated robust and similar nAb titers (*P*>0.05; **Figure 6e; Supplementary Table 2**). When comparing constructs at the low dose (0.6 μg), CprME/NS2B3 elicited higher nAb titers than the C2AprME construct (*P*<0.005). Titers were similar between mRNA-1893 and the other constructs when comparing within the 3- and 0.6-μg dose levels (*P*>0.05). Within each construct, both mRNA-1893 and C-2A-prME produced significant dose-dependent effects (*P*<0.01 and <0.001, respectively).

## Discussion

VLP-based vaccines offer several advantages compared with other modalities, such as monomeric proteins or whole-inactivated virus; these advantages include safety due to a lack of genetic material and an inability to replicate, and elicitation of strong immune responses due to native multi-array antigen presentation and efficient B-cell receptor engagement^37,38^. Delivery of VLPs by mRNA or other gene-based vectors adds the element of CD8 T cell induction not afforded by protein vaccines. ZIKV VLPs consisting of prME or CprME have been employed as the vaccine antigen across multiple vaccine platforms, including DNA, adenovirus-vectored, and purified particles^18,19,39–44^. Herein, we assessed the immune response elicited by mRNA-1325, which encodes the prME ORF from a Micronesia 2007 ZIKV isolate, and mRNA-1893, which encodes the contemporary RIO-U1 Zika prME ORF and differs from mRNA-1325 by 5 residues and the leader sequence.

This study suggested that the lower immunogenicity of mRNA-1325 compared with mRNA-1893 may arise from a defect in VLP production and secretion. In vitro characterization suggested that, although mRNA-1325 and mRNA-1893 expressed E protein intracellularly, E protein was only detectable in supernatant with transfection of mRNA-1893. Supernatant ELISAs and negative staining of purified particles, both using the VLP-specific antibody ZV-117, suggested that the secreted E protein generated by mRNA-1893 was in the form of a VLP. Protein half-life experiments demonstrated that E protein produced by mRNA-1325 has a shorter half-life than that produced by mRNA-1893 and is rapidly degraded by the proteasome. This rapid degradation may be due to partial misfolding, as the conformationally dependent antibody 4G2 did not detect E protein expression in cells transfected with mRNA-1325. While these in vitro defects may appear severe, the NHP data suggest that mRNA-1325 generates immunogens in vivo that elicit an antibody response.

The IgE_sp_-E(Ala40Thr) construct rescued VLP production to levels comparable to mRNA-1893 in vitro and generated nAb titers equivalent to mRNA-1893 in vaccinated mice, suggesting that the lower protection from ZIKV challenge afforded by mRNA-1325 compared with mRNA-1893 may be due to the Ala40Thr substitution in the E protein ORF. Of note, of the 1,509 ZIKV E protein sequences with data for position 40 in the Virus Pathogen Database and Analysis Resource (as of July 2022), most sequences contained Thr40 and only 1 sequence contained Ala40 (accession number EU545988); the latter sequence was used for the design of mRNA-1325. Ala40 may be detrimental to normal E protein function, explaining the lower protection afforded by mRNA-1325. Although it is unclear why this substitution would be deleterious to VLP production, this may be due to C-terminal serine endopeptidase cleavage between Lys41 and Pro42. When Pro is located at the C-terminal side of Lys, its ring structure restricts the freedom of rotation, making the polypeptide backbone more rigid. Lys-Pro bonds are typically cleavage-resistant but can be cleaved if the amino acid residues flanking the Lys-Pro sequence have small side chains that allow flexibility of the peptide bonds^45^. The Lys-Pro-Ala sequence of mRNA-1325 may make the Lys-Pro bond accessible to proteases, whereas in the Lys-Pro-Thr sequence of mRNA-1893, the bulky Thr side chain compared with Ala may prevent the torsion necessary for protease accessibility and cleavage.

Previous studies using a purified VLP vaccine platform reported that ZIKV C-containing VLPs are immunogenically superior to prME-only VLPs^39,40^. Here, we compared mRNA-1893 with 2 alternative designs that produce C-containing VLPs. The CprME ORF was encoded on a single mRNA, with cleavage of C from the downstream prME polyprotein achieved by the addition of a second mRNA encoding the NS2B3 protease or a 2A ribosome stop-go sequence between C and prME. Transfection of either CprME design resulted in production of secreted, C-containing VLPs that were detectable by the VLP-specific antibody ZV-117. Interestingly, although ELISAs conducted with neat supernatant did not reveal differences in the level of VLP production, CprME/NS2B3 transfection repeatedly resulted in a higher level of VLPs in any assay involving ultracentrifugation. Potential explanations for this observation include variation in particle size, centrifugation conditions, and/or the ability of particles to withstand purification. However, based exclusively on nAb titers, the inclusion of C in the VLP had a marginal effect on nAb titers in mice, as vaccination with equivalent doses of prME-only mRNA-1893 versus either of the 2 C-encoding mRNA constructs resulted in similar nAb titers; additional differences may bear out in challenge studies. These in vivo results, coupled with the observation that CprME-containing VLPs may be more resilient to ultracentrifugation, suggest that the differences between CprME and prME-only VLPs as immunogens may be influenced by vaccine platform. Of note, the addition of C to mRNA constructs may have practical implications, such as reduced efficiency in mRNA vaccine manufacturing due to the increased number of nucleotides^46^. Here, we provided evidence that mRNA-1893 is superior to mRNA-1325 in the rhesus macaque challenge model. Prime and boost vaccination with mRNA-1893 10 μg generated nAb titers comparable to those elicited by mRNA-1325 200 μg. In addition, no animals immunized with mRNA-1893 had detectable viral load at any time following challenge, whereas ≥ 1 animal in each of the mRNA-1325 dose groups exhibited viral breakthrough. Finally, no animals vaccinated with mRNA-1893 showed a boost in nAb titers following challenge, while ≥ 1 animal in each dose group of mRNA-1325 had a ≥ 4-fold increase in nAb titers at either 2 or 4 weeks following challenge, indicating induction of an anamnestic response. The lack of an anamnestic response and absence of detectable viremia in the animals receiving mRNA-1893 10 μg is indicative of rapid viral control. The viral breakthrough in animals vaccinated with mRNA-1325 despite comparable nAb titers between mRNA-1325 and mRNA-1893 could be due to differences in immune mediators like T-cell responses, antibody-dependent cell-mediated cytotoxicity, or antibody-dependent cellular phagocytosis that are not captured by neutralization assays. Indeed, previous reports have noted that antibody responses to DNA vaccine candidates expressing ZIKV prM-E are variably sensitive to prM content^47^; future studies should investigate antibodies against mature forms of the virion, as this may be more predictive than other neutralization assays.

In summary, results indicated that mRNA-1893 is superior to mRNA-1325. In addition to providing complete protection from ZIKV challenge in NHPs, mRNA-1893 generates higher nAb titers in mice than mRNA-1325 when given at an equivalent dose and elicits comparable titers at 1/20^th^ of the dose in NHPs. This enhanced immunogenicity may arise from a single amino acid residue difference in the E protein ORF of mRNA-1893, given that a Ala40Thr substitution rescues VLP secretion and restores immunogenicity of mRNA-1325 to levels comparable to mRNA-1893. Finally, CprME-encoding mRNA constructs do not outperform mRNA-1893 in mouse immunogenicity studies, suggesting that the prME-only VLPs produced by mRNA-1893 are not immunogenically inferior to C-containing VLPs. The data generated through preclinical animal models and in vitro characterization experiments support the advancement of mRNA-1893 into clinical trials. In a phase 1 trial (NCT04064905), mRNA-1893 had an acceptable safety profile and elicited strong anti-ZIKV serum nAb and binding antibody responses that persisted up to 1 year after vaccination. Results from a phase 1 trial (NCT03014089) showed that mRNA-1325 elicited poor immune responses; the preclinical data generated here may explain these clinical findings. Together, these preclinical and clinical results (reported elsewhere) support the continued development of mRNA-1893.

## Methods

### Generation of modified mRNA and LNPs

Modified mRNA was synthesized using in vitro T7 RNA polymerase-mediated transcription from a linearized DNA template, with N1-methyl-pseudouridine triphosphate and cap 1 to improve translation efficiency^48^. Formulations of mRNA-1325 and mRNA-1893 were prepared by dissolving ionizable lipids, distearoylphosphatidylcholine, cholesterol, and polyethylene glycol-lipid in ethanol (molar ratios of 50:10:38:5:1:5). This lipid mixture was combined with mRNA in a citrate buffer (50 mM, pH 4.0) at a ratio of 3:1 (aqueous:ethanol). The formulations were dialyzed against phosphate-buffered saline (pH 7.4) for <18 hours, concentrated, and passed through a 0.22-μm filter before storage at 4ºC. Final particle size and encapsulation were <100 nm and >80%, respectively, with endotoxin below 10 EU/mL^49^. The preclinical designations for mRNA-1325 and mRNA-1893 are CX-000171 and CX-005809, respectively; the studies described herein were conducted with preclinical material.

### Cell lines and antibodies

293T and Vero cells were obtained from the cell bank (Moderna, Inc.) and were cultured in Dulbecco’s modified Eagle’s medium supplemented with 10% fetal bovine serum. Human monoclonal antibodies (mAbs) ZV-116 and ZV-117 were provided by Dr. James Crowe (Vanderbilt University Medical Center, Nashville, TN). Mouse mAb ZV-67 was provided by Dr. Michael Diamond (Washington University School of Medicine, St. Louis, MO). Commercially available flavivirus E protein-specific mAb 4G2 (EMD Millipore, Darmstadt, Germany), rabbit anti-ZIKV prM and C antibodies (GeneTex, Irvine, CA), and goat horseradish peroxidase (HRP)-conjugated anti-human, anti-mouse, and anti-rabbit immunoglobulin G secondary antibodies (Southern Biotech, Birmingham, AL) were used.

### Collection of cell lysates and VLPs for use in Western blot, ELISA, and electron microscopy analyses

Using the TransIT-mRNA Transfection Kit (Mirus, Madison, WI), 1 × 10^7^ 293T cells were transiently transfected with mRNA (20 μg). Supernatant was collected 48 hours after transfection, centrifuged (500 x *g*) for 5 minutes, layered on a 25% glycerol/Tris/NaCl/EDTA cushion, and centrifuged (110,000 x *g*) for 2 hours at 4°C. Supernatants were aspirated and VLP-containing pellets were resuspended in phosphate-buffered saline. Isolated VLPs were analyzed by electron microscopy or Western blot. The corresponding 293T cells were lysed with RIPA buffer 48 hours after transfection. Protein concentrations were measured using a BCA assay kit (ThermoFisher). Western blot, ELISA, and electron microscopy experiments were performed using standard methodologies.

### Protein half-life experiments

293T cells (6.25 × 10^5^) were transfected with mRNA 1 μg for 24 hours, followed by treatment with CHX (100 μg/mL) ± chloroquine (50 μM) or MG132 (20 μM, Sigma) for varying amounts of time. Cells were lysed with RIPA buffer and blotted using ZV-116 or 4G2 antibodies. When 4G2 was used for Western blot, reducing reagent was omitted.

### Immunization studies

Experiments involving animals were carried out in compliance with approval from the Animal Care and Use Committee of Moderna, Inc. C57BL/6 mice (Charles River Laboratories, Wilmington, MA) were inoculated intramuscularly with 50 μL of mRNA/LNP vaccine on day 1; mice receiving 2 doses were immunized again on day 29. Blood, collected on days 28 and 56, was analyzed for serum nAbs.

Naïve Indian-origin rhesus macaques were inoculated intramuscularly with 500 μL of vaccine LNP constructs and challenged on day 56 with 1 × 10^3^ plaque-forming units ZIKV (strain PRVABC59). Blood was collected at time of challenge and at indicated time points after the challenge to measure viral load using quantitative reverse transcription polymerase chain reaction and serum nAb titers using the green fluorescent protein-expressing RVP neutralization assay^18,29^.

### Statistical methods

Sample sizes were determined based on the criteria set by the institutional Animal Care and Use Committee. One-way analysis of variance (ANOVA) was used to compare the effects of mRNA-1325 and mRNA-1893 and the different dosing regimens (1 vs 2 doses) on nAb titers in NHPs. This included testing of the dosing regimen as a single main effect and test article combinations. Two-way ANOVA was used to assess the dose response of CprME VLPs and prME-only VLPs and the effect of dose levels (3 vs 0.6 μg) on immunogenicity, including testing main effects of test articles and the dose levels and their interactions. Šidák correction was used to adjust for Type I error rate of multiple comparisons at an alpha level of 0.05, unless noted otherwise.

## Acknowledgments

Medical writing and editorial assistance were provided by Kate Russin, PhD, and Jessica Nepomuceno, PhD, of MEDiSTRAVA in accordance with Good Publication Practice (GPP3) guidelines, funded by Moderna, Inc., and under the direction of the authors. The authors would like to acknowledge James Crowe, MD (Vanderbilt University Medical Center) and Mike Diamond, MD, PhD (Washington University School of Medicine) for providing the antibodies used in this study; Arshan Nasir (Moderna, Inc.) for bioinformatics analysis of the sequence database; and Mario Roederer (NIAID, NIH) and Martha Nason (NIAID, NIH) for involvement in the NHP studies conducted by the Vaccinees Research Center at Bioqual. This work was supported in part by the Intramural Research Program of the NIAID. This project has been funded in whole or in part with federal funds from the Department of Health and Human Services; Office of the Assistant Secretary for Preparedness and Response; Biomedical Advanced Research and Development Authority, under Contract No. HHSO100201600029C.

## Author Contributions

BB, HB, SB, W-PK, BSG, KMM, TCP and AC contributed to the study concept and design. Data collection was performed by BB, NN, KEB, MA, TJR, KAD, and TCP; data analysis and interpretation was performed by BB, NN, KB, CJH, HB, BF, KEB, TJR, KAD, TCP, and AC. All authors contributed to the preparation of the manuscript and approved the final draft.

## Data Availability Statement

Upon request, and subject to review, Moderna, Inc., will provide the data that support the findings of this study.

