## Supplementary Table 1 and Supplementary Table 2 for "An optimized messenger RNA vaccine candidate protects non-human primates from Zika virus infection"

**Supplementary Table 1. Summary of statistical comparisons between nAb titers measured at week 4 (day 28) and week 8 (day 56) in NHPs**

| Comparisons | Adjusted *P* Values | |
|---|---|---|
| | **Week 4** | **Week 8** |
| Control, 200 µg x2 vs. mRNA-1325, 10 µg x2 | 0.0059 | <0.0001 |
| Control, 200 µg x2 vs. mRNA-1325, 50 µg x2 | 0.0017 | 0.0004 |
| Control, 200 µg x2 vs. mRNA-1325, 200 µg x1 | 0.0004 | 0.0025 |
| Control, 200 µg x2 vs. mRNA-1325, 200 µg x2 | 0.0002 | <0.0001 |
| Control, 200 µg x2 vs. mRNA-1893, 10 µg x2 | 0.0024 | <0.0001 |
| mRNA-1325, 10 µg x2 vs. mRNA-1325, 50 µg x2 | 0.3309 | 0.4673 |
| mRNA-1325, 10 µg x2 vs. mRNA-1325, 200 µg x1 | 0.0214 | 0.9571 |
| mRNA-1325, 10 µg x2 vs. mRNA-1325, 200 µg x2 | 0.0511 | 0.0007 |
| mRNA-1325, 10 µg x2 vs. mRNA-1893, 10 µg x2 | 0.0851 | 0.0459 |
| mRNA-1325, 50 µg x2 vs. mRNA-1325, 200 µg x1 | 0.4120 | 0.2916 |
| mRNA-1325, 50 µg x2 vs. mRNA-1325, 200 µg x2 | 0.9795 | 0.4697 |
| mRNA-1325, 50 µg x2 vs. mRNA-1893, 10 µg x2 | 0.8869 | 0.9600 |
| mRNA-1325, 200 µg x1 vs. mRNA-1325, 200 µg x2 | 0.8678 | 0.0379 |
| mRNA-1325, 200 µg x1 vs. mRNA-1893, 10 µg x2 | 0.9995 | 0.0568 |
| mRNA-1325, 200 µg x2 vs. mRNA-1893, 10 µg x2 | >0.9999 | 0.9719 |

nAb, neutralizing antibody; NHP, non-human primate.

**Supplementary Table 2. Summary of statistical comparisons between nAb titers measured on day 56 in mice**

| Comparisons | Adjusted *P* Values |
|---|---|
| 2+1µg:mRNA-1893+Control vs. 2+1µg:CprME+NS2B3 | 0.9988 |
| 2+1µg:mRNA-1893+Control vs. 2+1µg:C2AprME+Control | 0.9810 |
| 2+1µg:mRNA-1893+Control vs. 0.4+0.2µg:mRNA-1893+Control | 0.0036 |
| 2+1µg:mRNA-1893+Control vs. 0.4+0.2µg:CprME+NS2B3 | 0.6693 |
| 2+1µg:mRNA-1893+Control vs. 0.4+0.2µg:C2AprME+Control | <0.0001 |
| 2+1µg:CprME+NS2B3 vs. 2+1µg:C2AprME+Control | 0.4435 |
| 2+1µg:CprME+NS2B3 vs. 0.4+0.2µg:mRNA-1893+Control | 0.0002 |
| 2+1µg:CprME+NS2B3 vs. 0.4+0.2µg:CprME+NS2B3 | 0.1155 |
| 2+1µg:CprME+NS2B3 vs. 0.4+0.2µg:C2AprME+Control | <0.0001 |
| 2+1µg:C2AprME+Control vs. 0.4+0.2µg:mRNA-1893+Control | 0.1098 |
| 2+1µg:C2AprME+Control vs. 0.4+0.2µg:CprME+NS2B3 | >0.9999 |
| 2+1µg:C2AprME+Control vs. 0.4+0.2µg:C2AprME+Control | 0.0006 |
| 0.4+0.2µg:mRNA-1893+Control vs. 0.4+0.2µg:CprME+NS2B3 | 0.4283 |
| 0.4+0.2µg:mRNA-1893+Control vs. 0.4+0.2µg:C2AprME+Control | 0.7084 |
| 0.4+0.2µg:CprME+NS2B3 vs. 0.4+0.2µg:C2AprME+Control | 0.0043 |

CprME, capsid, premembrane/membrane, and envelope; nAb, neutralizing antibody.
